# methylSCOPA and META-methylSCOPA: software for the analysis and aggregation of epigenome-wide association studies of multiple correlated phenotypes

**DOI:** 10.1101/656918

**Authors:** Harmen Draisma, Jun Liu, Igor Pupko, Ayşe Demirkan, Zhanna Balkhiyarova, Andrew P Morris, Reedik Magi, Matthias Wielscher, Saqib Hassan, Cornelia M. van Duijn, Sylvain Sebert, Marjo-Riitta Jarvelin, Marika Kaakinen, Inga Prokopenko

## Abstract

**Background:** Multi-phenotype genome-wide association studies (MP-GWAS) of correlated traits have greater power to detect genotype–phenotype associations than single-trait GWAS. However, no multi-phenotype analysis method exists for epigenome-wide association studies (EWAS).

**Results:** We extended the SCOPA approach developed by us to “methylSCOPA” software in C++ by ‘reversely’ regressing DNA hyper/hypo-methylation information on a linear combination of phenotypes. We evaluated two models of association between DNA methylation and fasting glucose (FG) and insulin (FI) levels: Model 1, including FG, FI, and three measured potential confounders (body mass index [BMI], fasting serum triglyceride levels [TG], and waist/hip ratio [WHR]), and Model 2, including FG and FI corrected for the effects of BMI, TG, and WHR. Both models were additionally corrected for participant sex and smoking status (current/ever/never). We meta-analyzed the cohort-specific MP-EWAS results with our novel software META-methylSCOPA, mapped genomic locations to CGCh37/hg19, and adopted *P*<1×10^−7^ to denote epigenome-wide significance. We used the Illumina Infinium HumanMethylation450K BeadChip array data from the Northern Finland Birth Cohorts (NFBC) 1966/1986. We quality-controlled the data, regressed out the effects of measured potential confounders, and normalized the methylation signal intensity and FI data. The MP-EWAS included data for 643/457 individuals from NFBC1966 and NFBC1986, respectively (total N=1,100).

In Model 1, we detected epigenome-wide significant association in the MP-EWAS meta-analysis at cg13708645 (chr12:121,974,305; *P*=1.2×10^−8^) within *KDM2B* gene. Single-trait effects within *KDM2B* were on FI, BMI, and WHR. Model with effect on BMI and WHR showed the strongest association at this locus, while effect on FI in single-phenotype analysis was driven by the effect of adiposity. In Model 2, the strongest association was at cg05063096 (chr3:143,689,810; *P*=2.3×10^−7^) annotated to *C3orf58* with strongest effect on FI in single-trait analysis and multi-phenotype effect on FI and WHI within Model 1.

We characterized the effects of established EWAS loci for diabetes and its risk factors and detected suggestive (p<0.01) associations at six markers including *PHGDH, TXNIP, SLC7A11, CPT1A, MYO5C* and *ABCG1*, through the dissection of the multi-phenotype effects in Model 1.

**Conclusions:** We implemented MP-EWAS in methylSCOPA and demonstrated its enhanced power over single-trait EWAS for correlated phenotypes in large-scale data.

## Background

Multi-phenotype genome-wide association studies (MP-GWAS) of correlated traits are more powerful, give better precision of estimates, and provide enhanced biological insight, i.e. suggestion of potential pleiotropic effects, as compared to single-phenotype GWAS^1–6^. We have previously developed an MP-GWAS method using the “reverse regression” approach in which allele dosage is regressed on a linear combination of phenotypes, implemented into the software tool SCOPA and meta-analysis tool METASCOPA^7^. However, no multi-phenotype epigenome-wide association study (MP-EWAS) method exists, although EWAS have recently gained increased attention due to advances in technology and thus lowered costs of measuring epigenetic regulation.

DNA methylation is a type of epigenetic regulation and is most widely used within EWAS. Methylation refers to the attachment of methylation groups to the DNA molecule. Methylation of CpG islands within a gene’s promoter usually implies that that the gene is not transcribed. DNA methylation is tissue-specific, reversible, and inheritable. Usually, the cytosine copies on both strands are either methylated or unmethylated.

In correlated traits, there is a considerable, although incomplete, overlap between the measures of glucose homeostasis and type 2 diabetes (T2D). For instance, the genetic correlation between fasting glucose levels (FG) and fasting insulin levels (FI) estimated by cross-trait LD Score regression is 0.31, and the genetic correlations between FG and T2D and between FI and T2D are 0.58 and 0.48, respectively^8^. The study of glycaemic traits in healthy individuals can provide insights about the pathophysiology of T2D, and the (epi)genetic study on these phenotypes can inform on the molecular mechanisms leading to T2D – also those influenced by individual’s lifestyle and environmental exposures, as they have been shown to leave a mark on the individual’s epigenome^9^. One of the advantages of studying glycaemic traits, rather than T2D, is that sample sizes can be much larger, as they are independent of T2D prevalence. Indeed, genome-wide methylation in blood has been associated with body mass index, T2D and measures of glucose metabolism^10–12^. However, no study has previously aimed at unravelling the epigenetics of these traits by taking into account their correlations with each other, most likely due to the lack of appropriate methodology.

Our aims in the current work were two-fold. First, we aimed to extend the reverse regression approach for methylation data and implement it in a software tool. We addressed this aim by developing methylSCOPA (Software for COrrelated Phenotype Analysis with methylation data), which is the SCOPA extension for DNA methylation data. methylSCOPA association summary statistics can also be aggregated across EWAS through fixed-effects meta-analysis, implemented in META-methylSCOPA, which is the META-SCOPA extension for MP-EWAS meta-analysis. Analogous to META-SCOPA, this enables application of reverse regression in large-scale international consortia efforts where, for instance, ethical concerns and legal restrictions preclude joint analysis of individual-level genome-wide DNA methylation and phenotype data from different studies. Second, we aimed to test the method for epigenetic effects on FG & FI variability. We report one novel methylation probe associated with FG and FI from these analyses and dissect the multi-phenotype epigenetic effects at 11 established methylation marks for metabolic traits.

## Implementation

### Reverse regression model of multiple correlated phenotypes

methylSCOPA extends the SCOPA analysis framework^7^ to the analysis of DNA methylation data. Specifically, this means that methylSCOPA allows for methylation (instead of genotype as in SCOPA) data as input, and that it analyses methylation data analogous to the way in which SCOPA analyses genotype dosage data.

DNA methylation assays in tissue samples return at any given site an average methylation percentage for a mixture of cells. These percent methylation values are continuous and range from 0 to 100^13^. In methylSCOPA we model these percent methylation values as a function of the observed phenotypes using linear reverse regression, analogous to how SCOPA models the genotype at a single-nucleotide polymorphism (SNP) as a function of the observed phenotypes. Therefore, analogous to expression (1) from Mägi *et al.*^7^, considering a sample of unrelated individuals with *J* phenotypes denoted by *y*_1_, *y*_2_, …, *y*_*J*_, in methylSCOPA we model the DNA methylation value *Methylation*_*i*_ at a particular probe for individual *i* as

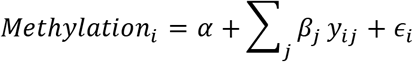

In this expression, *y*_*ij*_ denotes the phenotypic value for individual *i*, phenotype *j*; *β*_*j*_ denotes the effect of the *j*th phenotype on the degree of DNA hyper/hypomethylation at the probe (analogous to the effect *β*_*j*_ of the *j*th phenotype on genotype at the SNP under consideration in the SCOPA model); and ∈_*i*_ ~*N*(0, *σ*^2^), where *σ*^2^ is the residual variance. We recommend that covariates relevant to the multi-phenotype effects should be included in the model; otherwise, confounding factors should be regressed out of the phenotypes and resulting residuals should be used instead.

For further dissection of epigenome-wide significant (*p*<1×10^−7^) multiple-phenotype association signals, the analysis of different phenotype combinations is enabled. We assess the model fit within each phenotype combination through the use of the Bayesian Information Criterion (BIC), with the smallest value indicating the best fit.

Meta-analysis of multiple EWAS of the same set of correlated phenotypes is enabled through the application of the method for the synthesis of regression slopes^14^, similarly to METASCOPA^7^. We further implemented model selection for the signals reaching epigenome-wide significance in the meta-analysis by using the ‘meta-Bayesian Information Criterion’ (meta-BIC) value. Following the notation from Bohning^15^, the likelihood is defined as

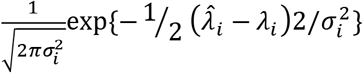

and the meta-likelihood as

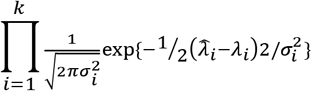

The value of the Bayesian Information Criterion is calculated as in Wit *et al.*^16^

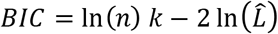

where

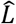 = maximized value of likelihood function
*n* = (combined) sample size
*k* = number of parameters estimated by the model

Therefore, the value of the meta-log-likelihood can simply be computed as the sum of the log-likelihoods of the individual studies, and the ‘meta-BIC’ value for the meta-analysis result can subsequently be computed based on that sum, i.e., based on the meta-log-likelihood.

### methylSCOPA and META-methylSCOPA

As methylSCOPA and META-methylSCOPA are the extensions for methylation data analysis of SCOPA and META-SCOPA^7^, respectively, installation procedures, command line options, and output columns for methylSCOPA and META-methylSCOPA are largely analogous to those for SCOPA and META-SCOPA, respectively and are detailed in the tutorial provided along the software.

### Required file formats

Similarly to SCOPA^7^, methylSCOPA requires phenotype data in GEN/SAMPLE format utilized by the IMPUTE and SNPTESTv.2. The methylation data file format required by the software is described in the tutorial available online http://www.imperial.ac.uk/people/h.draisma/research.html.

## Case study

### Study populations

To implement our novel methylSCOPA and META-methylSCOPA methods and tools, we performed an MP-EWAS of two correlated glycaemic traits: FI and FG in two independent cohorts, followed by their meta-analysis. These two independent cohorts are the Northern Finland Birth Cohorts (NFBC) 1966/1986 which cover almost all births in the two northernmost provinces of Finland between the expected dates of delivery falling in 1966^17^ (N=12,058 live-born children) and between 1^st^ of July 1985 and 30^th^ June 1986^18^ (N=9,432 live-born children). The children born to the cohort have been followed up throughout their lives, and here, we used data from the 31-year clinical examination for NFBC1966 and from the 15/16 year clinical examination for the NFBC1986. FI and FG were measured after overnight fasting and processed according to the standard protocol. The methylation data were obtained from the Illumina Infinium HumanMethylation450K BeadChip array for 807 randomly selected subjects that had provided fasting blood samples at the 31-year clinical examination (NFBC1966) and for 15-16-year-old individuals (NFBC1986). After quality control, the MP-EWAS included data for 643/457 individuals from NFBC1966 and NFBC1986, respectively. All the individuals included in the study have provided written informed consent (or parents of the participants of NFBC1986). The study was approved by the ethical committees of the University of Oulu and Imperial College London (Approval:18IC4421).

### Quality control of the methylation data

Methylation data was quality controlled as follows: we 1) removed duplicate samples, 2) filtered based on methylation detection *P*-value, 3) performed subset quantile normalization of raw methylation signal intensity values^19^, 4) removed methylation data batch effects using the pipeline CPACOR^20^, 5) detected and removed methylation data outliers as well as samples with gender mismatch, 6) applied white blood cell type composition correction, using “Houseman estimates”^21^, 7) transformed the values resulting from the previous step (step 6) to “beta” values, 8) applied inverse normal transformation to the methylation “beta” values, and finally 9) calculated residuals of the inverse normal-transformed methylation “beta” values by linear regression, using as covariates the Houseman estimates, first 30 control probe PC scores, participant sex, smoking status (current/ever/never as defined in keeping with the definitions as used for NFBC1966 and NFBC1986 within the CARTA consortium [Morris et al., BMJ Open 2015;5:e008808], and also including an additional category ‘unknown’ for participants with missing data on smoking status), and – in the case of Model 2 analysis – BMI, TG, and WHR.

### Phenotype definitions, imputation and transformations

We used fasting plasma glucose values (mmol/l) and fasting circulating insulin values (pmol/l) as the main variables of interest in our MP-EWAS. FG was used as such whereas we applied natural log-transformation to FI. We excluded from the analyses non-fasting individuals, pregnant women, diabetic individuals defined as fasting plasma glucose>=7mmol/l; 2-hour post oral glucose tolerance test glucose>=11.1mmol/L, HbA1c>=6.5%; diagnosis of type 1 or type 2 diabetes; or on diabetes treatment [oral and insulin] and individuals on lipid-lowering medication (ATC code “A10”). We then imputed the phenotype data for males and females separately with random forest (MisForest in R). We included use of oral contraceptives for the imputation model for females and additionally for both we included other available indicators of metabolic health, such as blood pressure and metabolomic data. The data for both sexes were combined after imputation. Finally, we regressed out participant sex and smoking status (current/ever/never/unknown), and – in the case of Model 2, see below – BMI, TG, and WHR from the FI and FG values. In fixed-effects meta-analysis of NFBC1966 and NFBC1986 using the functions “escalc” and “rma” in the “R” package ‘metafor’, the Pearson’s product-moment correlation between the resulting FG and FI residuals used for Model 2 analysis was 0.23 (*P*<0.0001).

### MP-EWAS

We fitted two models: Model 1 allows for the detection of effects that might be shared across ‘phenotypes of interest’ (FG and FI in this case) and measured potential confounders, i.e. confounders were included in the same model. The confounders considered were triglycerides (TG), body mass index (BMI) and waist/hip ratio (WHR). We regressed out the effect of sex and smoking status (current/ever/never/unknown) from these confounders, similarly as we did for the measurements of FG and FI. Model 2 includes multi-phenotype EWAS of FI/FG corrected for measured potential confounders and allows for the detection of effects that are unique to the phenotypes of interest, i.e. which are not shared with the measured potential confounders. In other words, Model 2 had the measured potential confounders TG, BMI and WHR regressed out from the FG and FI values prior to model fitting. Mathematically, the two MP-EWAS models fitted were as follows:

1. Methylation probe “beta” residuals value_i_ = β_0_ + β_1_FGres_i_ + β_2_ln(FI)res_i_ + β_3_BMIres + β_4_TGres + β_5_WHRres+ e_i_
2. Methylation probe “beta” residuals value_i_ = β_0_ + β_1_FGres_i_ + β_2_ln(FI)res_i_ + e_i_

where i=1,…,*n*, FGres = fasting glucose residuals, ln(FI)res = residuals of natural log-transformed fasting insulin, BMIres = body mass index, TGres = fasting serum triglyceride level and WHRres = waist/hip ratio.

We meta-analyzed the cohort-specific MP-EWAS results with META-methylSCOPA, mapped genomic locations to CGCh37/hg19, and adopted *P*<1×10^−7^ to denote epigenome-wide significance. From the META-methylSCOPA meta-analysis results, we filtered out associations involving probes that could be cross-reactive, polymorphic, or have been suggested to be excluded from analysis for any other reason by Chen et al. [Chen et al., Epigenetics. 2013 Jan 11;8(2)] (their files “List of cross-reactive probes” and “List of polymorphic CpGs”), Naeem et al. [Naeem et al., 2017; Zhou et al., 2017] (column “MASK.general” as in their file “hm450.hg19.manifest.tsv.gz” and columns “MASK_general_FIN” and “MASK_general_EUR” as in their file “hm450.hg19.manifest.pop.tsv.gz” as on “http://zwdzwd.github.io/InfiniumAnnotation#download” [accessed 2019-05-19 BST]).

## Results and discussion

Our MP-EWAS (**Figure 1**) of 466,342 methylation probes and FG and FI using methylSCOPA and META-methylSCOPA resulted in one epigenome-wide significant signal at cg13708645 (chr12:121,974,305; *P_Model1*=1.2×10^−8^; *P_Model2*=0.48) within *KDM2B* (**Table 1**, **Figure 1**), and one at suggestive (*P*<10^−6^) significance at cg05063096 (chr3:143,689,810; *P_Model1*=2.0×10^−7^; *P_Model2*=2.3×10^−7^) within *C3orf58* (**Table 1, Figure 2**).

**Table 1.** Lead cgIDs in META-methylSCOPA meta-analysis of fasting glucose (FG) and fasting insulin (FI) in 1,100 individuals from NFBC1966 and NFBC1986 in Models 1 and 2. Measured potential confounders body mass index (BMI), triglycerides (TG) and waist-to-hip ratio (WHR) were included in Model 1 whereas their effects were regressed out in Model 2.

| Model | Locus | Lead cgID | Chr | Position<sup>a</sup> (bp) | FG effect<br>(SE) | FI effect<br>(SE) | BMI effect<br>(SE) | TG effect<br>(SE) | WHR effect<br>(SE) | P-value |
| --- | --- | --- | --- | --- | --- | --- | --- | --- | --- | --- |
| **1** | **KDM2B** | **cg13708645** | **12** | **121,974,305** | **-0.002**<br>**(0.003)** | **0.004**<br>**(0.004)** | **0.001**<br>**(4.6×10<sup>-4</sup>)** | **-4.7×10<sup>-4</sup>**<br>**(0.002)** | **0.075**<br>**(0.027)** | **1.2×10<sup>-8</sup>** |
| 1 | *C3orf58* | cg05063096 | 3 | 143,689,810 | 0.002<br>(7.9×10<sup>-4</sup>) | -0.006<br>(0.001) | 7.0×10<sup>-5</sup><br>(1.2×10<sup>-4</sup>) | 0.002<br>(6.4×10<sup>-4</sup>) | 0.022<br>(0.007) | 2.0×10<sup>-7</sup> |
| 2 | *KDM2B* | cg13708645 | 12 | 121,974,305 | -0.002<br>(0.003) | 0.005<br>(0.004) |  |  |  | 0.48 |
| **2** | ***C3orf58*** | **cg05063096** | **3** | **143,689,810** | **0.002**<br>**(7.9×10<sup>-4</sup>)** | **-0.006**<br>**(0.001)** |  |  |  | **2.3×10<sup>-7</sup>** |

Chr: chromosome. SE: standard error. FG: fasting plasma glucose level. FI: fasting circulating insulin level. BMI: body mass index. TG: fasting serum triglyceride level. WHR: waist-to-hip ratio
<sup>a</sup>Position reported for NCBI build GRCh37 (UCSC hg19 assembly)

**Figure 1.**
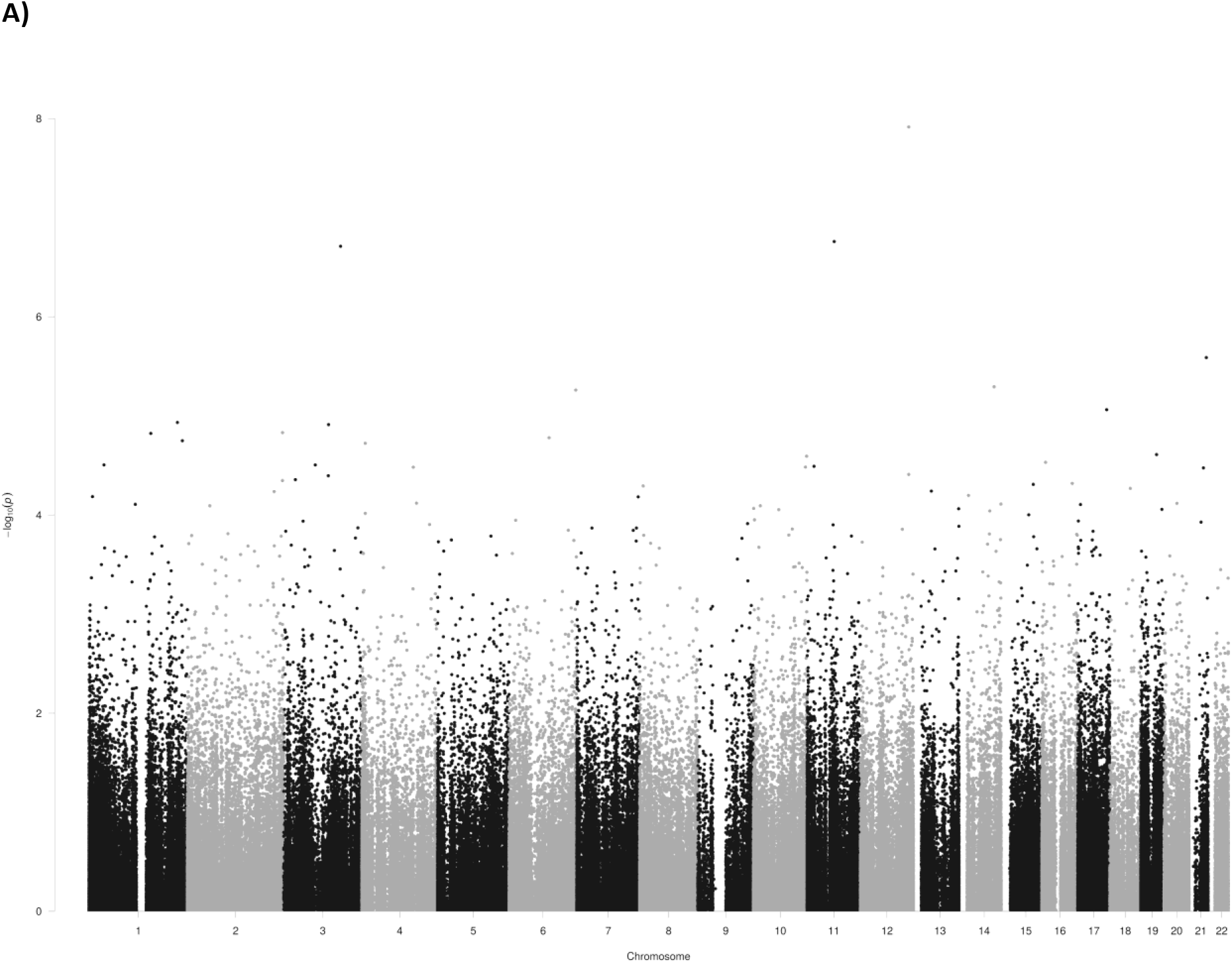
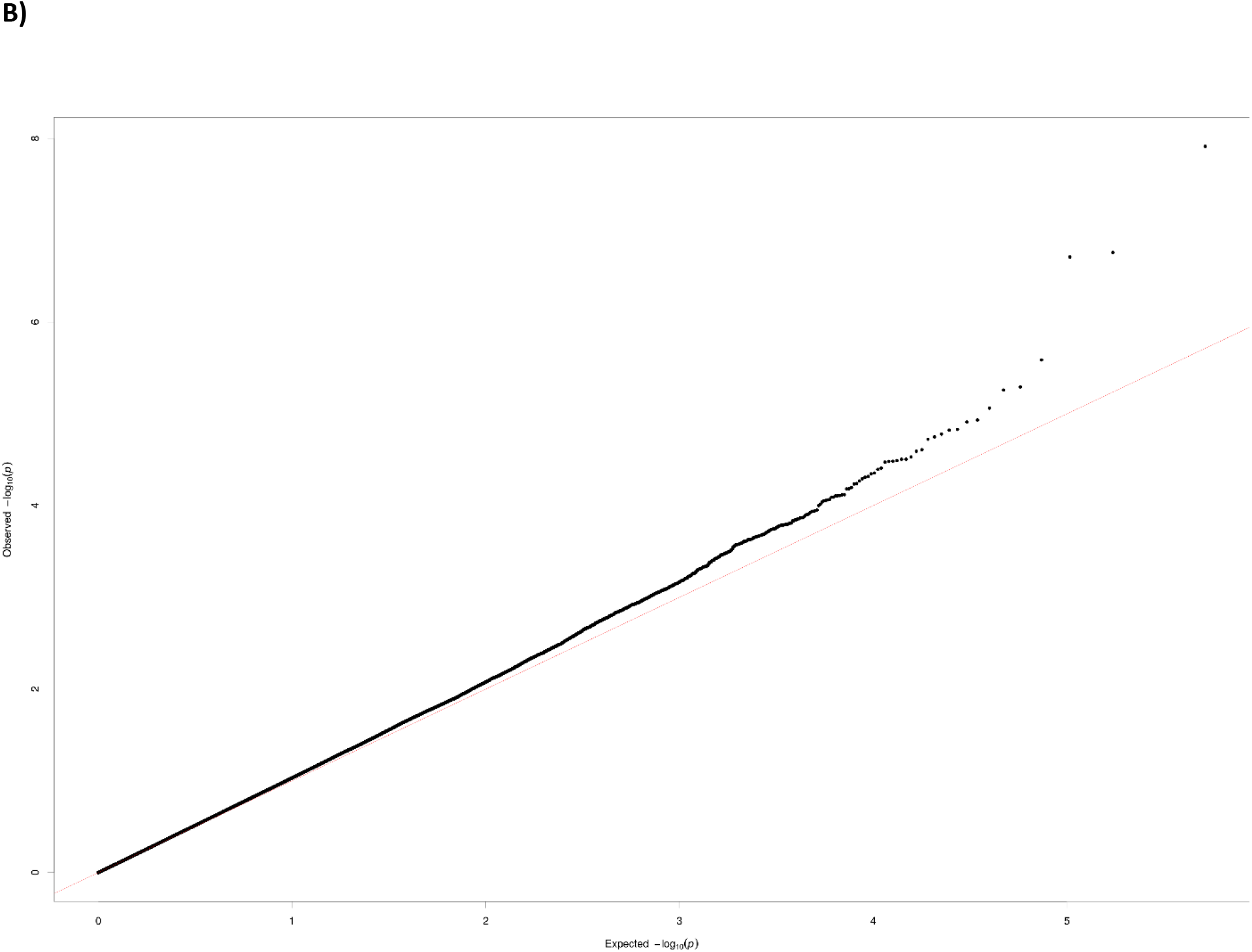
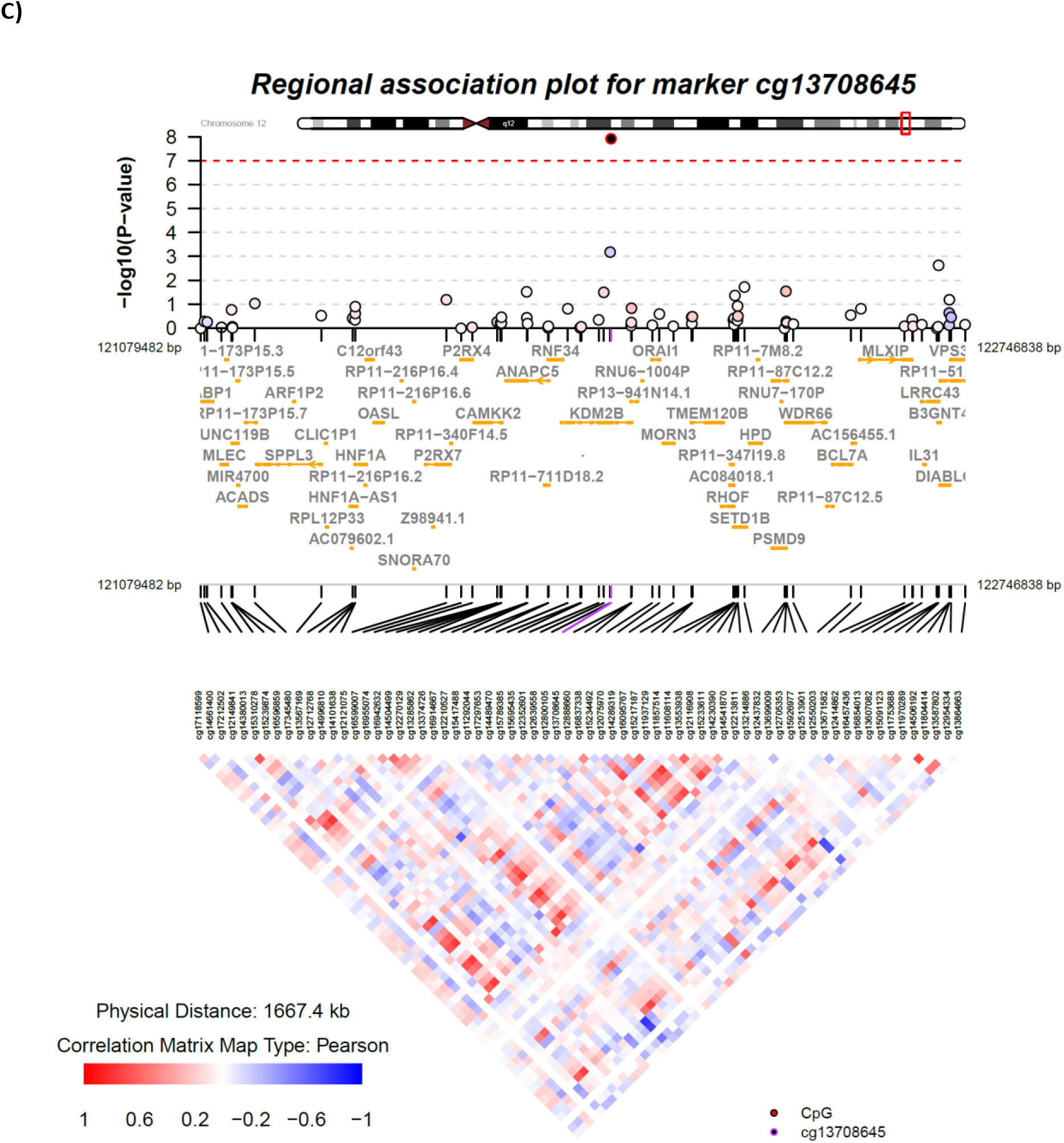
Results from the meta-analysis of MP-EWAS for FG and FI from Model 1. A) Manhattan plot, B) QQ-plot, and C) Regional association plot for the top signal. In the regional association plot, each point represents a CpG passing quality control in the association analysis, plotted with their *p*-value (on a −log_10_ scale) as a function of genomic position (NCBI build GRCh37, UCSC hg 19 assembly). The lead CpG is represented by the circle with the red edge and black face. The color coding of all other CpGs indicates Pearson correlation with the lead CpG in meta-analysis of NFBC1966 age 31 and NFBC1986 data. We performed the meta-analysis of the correlations between CpGs using function “escalc” from the ‘metafor’ package in the statistical language and environment “R” [R package], and created the signal plots using the coMET package. Gene annotations are taken from the University of California Santa Cruz genome browser

**Figure 2.**
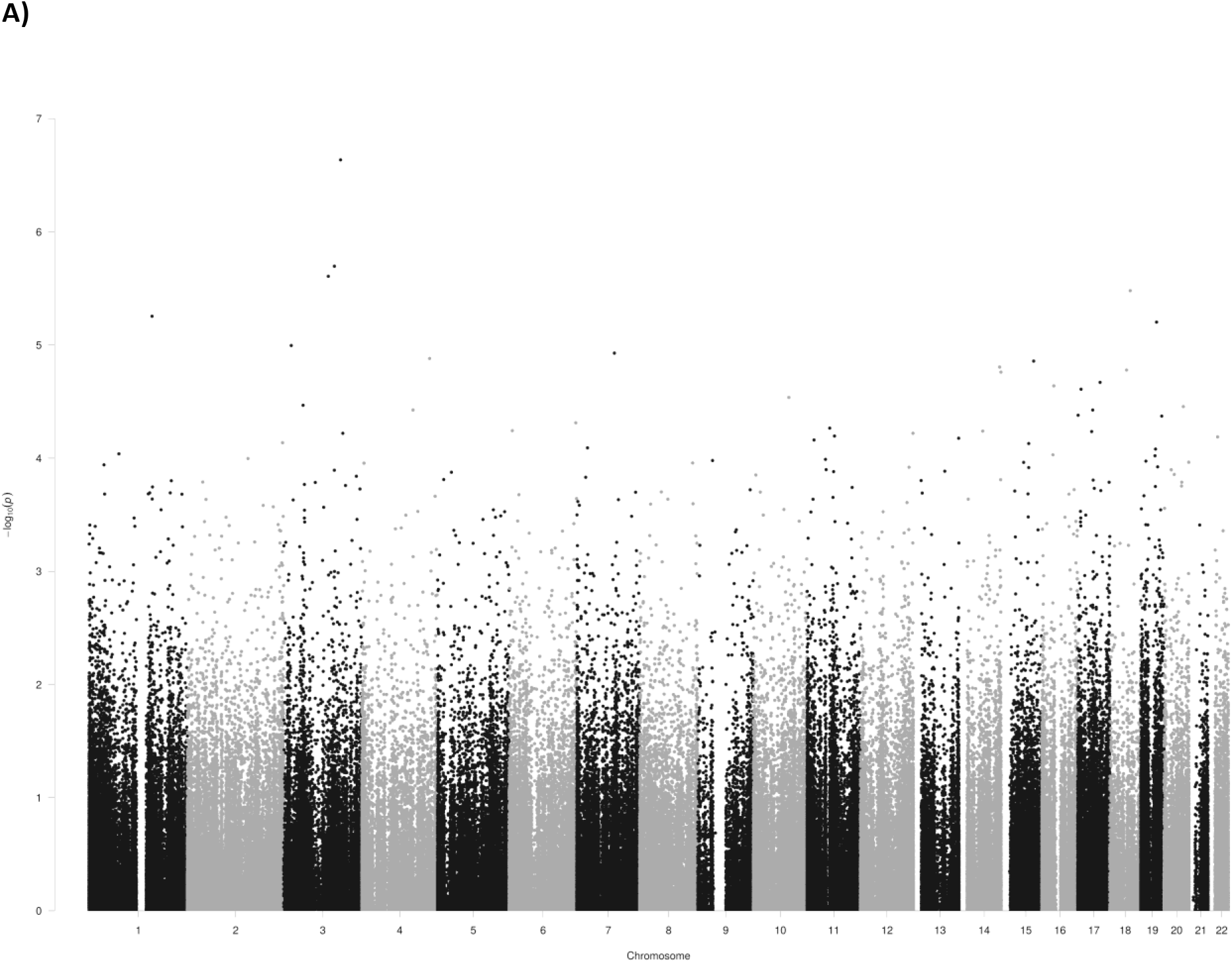
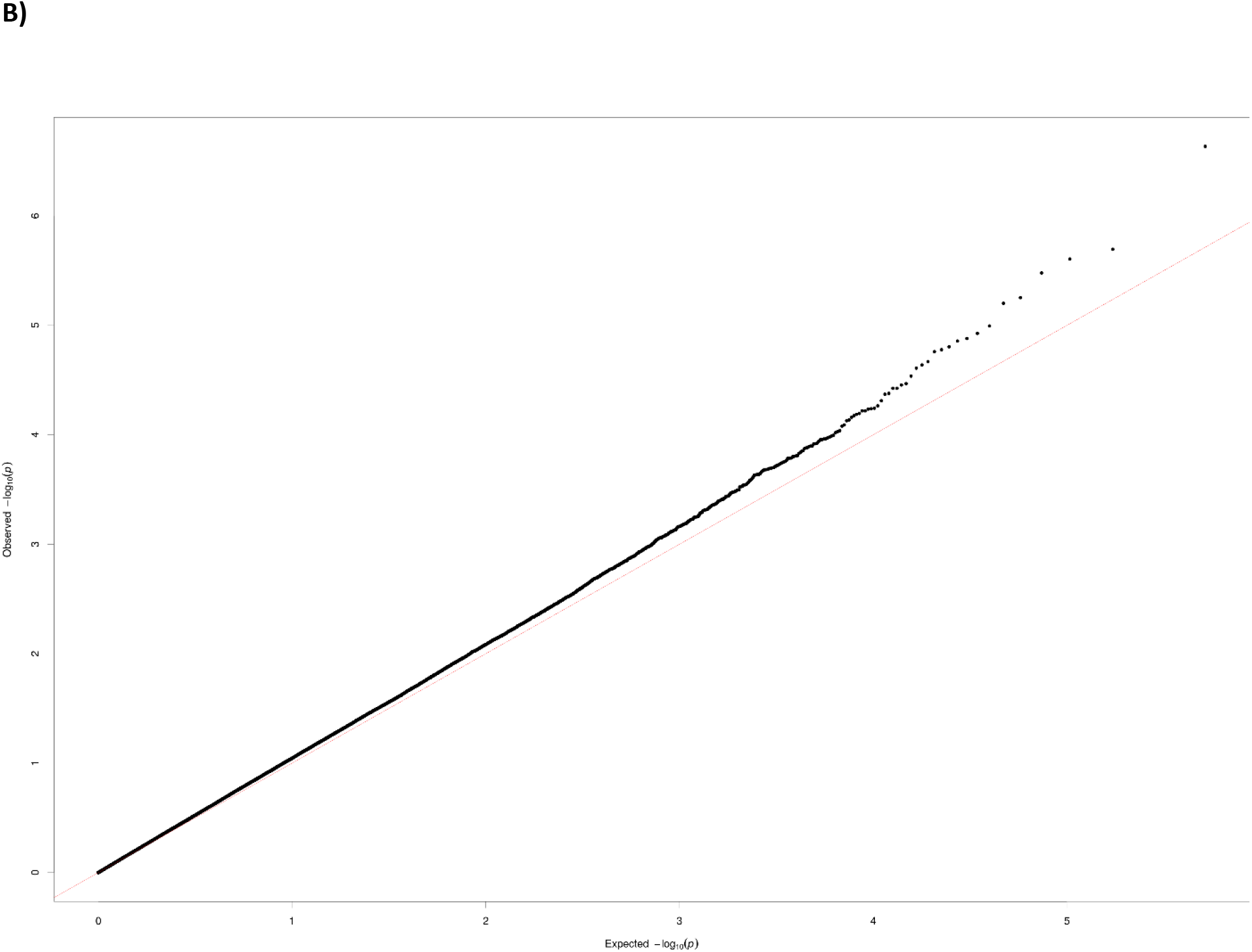
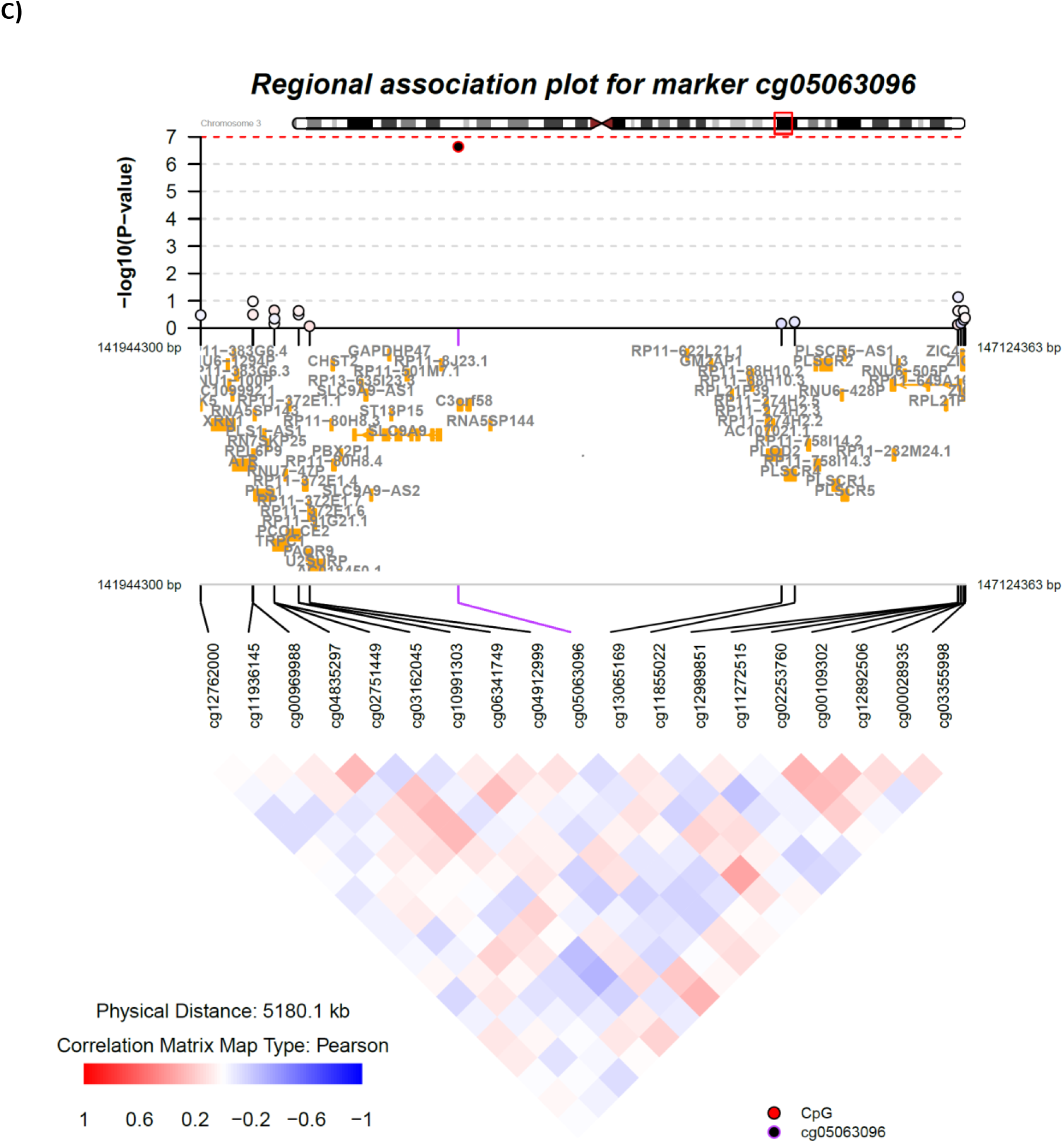
Results from the meta-analysis of MP-EWAS for FG and FI from Model 2. A) Manhattan plot, B) QQ-plot, and C) Regional association plot for the top signal. In the regional association plot, each point represents a CpG passing quality control in the association analysis, plotted with their *p*-value (on a −log_10_ scale) as a function of genomic position (NCBI build GRCh37, UCSC hg 19 assembly). The lead CpG is represented by the circle with the red edge and black face. The color coding of all other CpGs indicates Pearson correlation with the lead CpG in meta-analysis of NFBC1966 age 31 and NFBC1986 data. We performed the meta-analysis of the correlations between CpGs using function “escalc” from the ‘metafor’ package in the statistical language and environment “R” [R package], and created the signal plots using the coMET package. Gene annotations are taken from the University of California Santa Cruz genome browser

The locus at *KDM2B* has the most pronounced association with FI, BMI and WHR in univariate analyses (**Table 2**). It was not significant in Model 2. Model selection for this locus though metaBIC identified BMI and WHR as the most significant phenotype combination (**Table 3**), the effecton FI at this locus was driven through its effect on obesity phenotypes. The locus at *C3orf58*, identified within Model 2 with suggestive significance, showed stronger effect (**Table 1**), when additional phenotypes were considered in Model 1. The strongest phenotype effects at *C3orf58* were on combination of FI and WHR, while the effect on FI in univariate analysis would have not led to the detection of this efpigenetic effect.

**Table 2.**
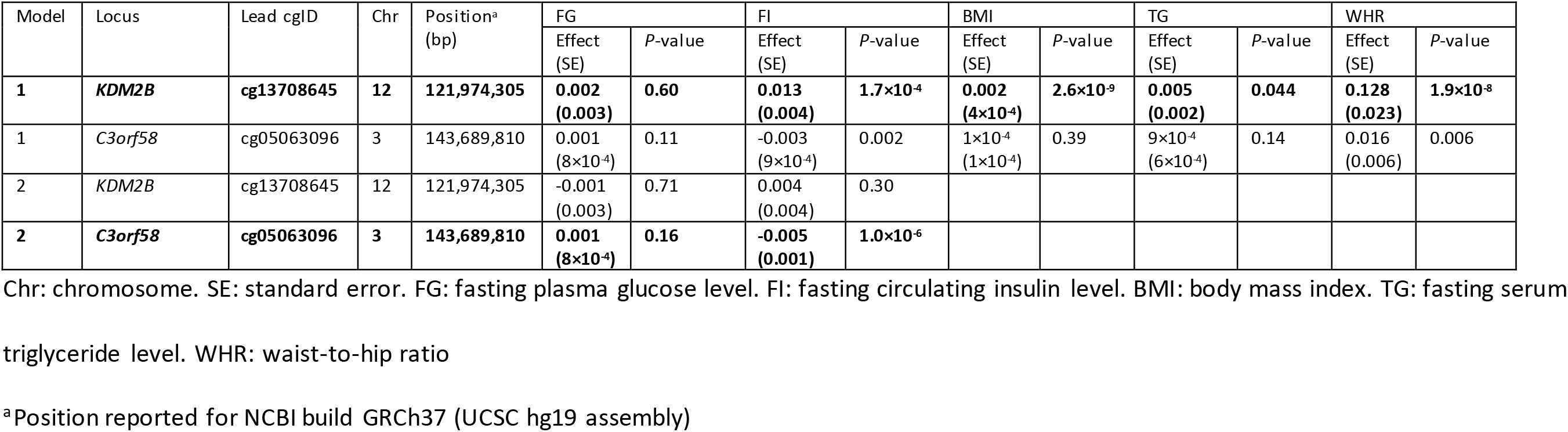
Univariate EWAS meta-analysis at lead cgIDs of FI and FG in 1,100 individuals from NFBC1966 and NFBC1986. Measured potential confounders body mass index (BMI), triglycerides (TG) and waist-to-hip ratio (WHR) were included in Model 1 whereas their effects were regressed out in Model 2.

**Table 3.** Dissection of the multiple phenotype association signals for the lead cgIDs in the MP-EWAS of fasting glucose (FG) and fasting insulin (FI) in 1,100 individuals from NFBC1966 and NFBC1986 in Models 1 and 2. The values displayed are the differences in metaBIC from that of the null model. Potential confounders of body mass index (BMI), triglycerides (TG) and waist-to-hip ratio (WHR) were included in Model 1 whereas their effects were regressed out in Model 2.

| Model | Model 1<br>cg13708645 | Model 1<br>cg05063096 | Model 2<br>cg13708645 | Model 2<br>cg05063096 |
| --- | --- | --- | --- | --- |
| BMI | -28.26 | 6.10 |  |  |
| BMI+FG | -26.33 | 10.59 |  |  |
| BMI+FG+FI | -21.73 | -2.57 |  |  |
| BMI+FG+TG | -21.23 | 15.91 |  |  |
| BMI+FG+TG+FI | -16.57 | -2.63 |  |  |
| BMI+FG+TG+WHR | -23.45 | 14.77 |  |  |
| BMI+FG+TG+WHR+FI | -17.88 | -8.40 |  |  |
| BMI+FG+WHR | -28.67 | 8.95 |  |  |
| BMI+FG+WHR+FI | -23.11 | -8.85 |  |  |
| BMI+FI | -23.87 | -4.17 |  |  |
| BMI+TG | -23.21 | 11.51 |  |  |
| BMI+TG+FI | -18.35 | -3.33 |  |  |
| BMI+TG+WHR | -25.69 | 10.29 |  |  |
| BMI+TG+WHR+FI | -20.15 | -8.57 |  |  |
| BMI+WHR | -30.83 | 4.43 |  |  |
| BMI+WHR+FI | -25.69 | -9.87 |  |  |
| FG | 0.18 | 3.98 | 1.70 | 4.68 |
| FG+FI | -8.47 | -3.53 | 7.09 | -18.33 |
| FG+TG | 2.47 | 8.97 |  |  |
| FG+TG+FI | -2.85 | -5.31 |  |  |
| FG+TG+WHR | -19.33 | 9.73 |  |  |
| FG+TG+WHR+FI | -16.97 | -15.23 |  |  |
| FG+WHR | -24.01 | 3.65 |  |  |
| FG+WHR+FI | -22.47 | -15.55 |  |  |
| FI | -9.72 | -4.96 | 5.66 | -18.90 |
| TG | 1.98 | 4.78 |  |  |
| TG+FI | -3.75 | -5.81 |  |  |
| TG+WHR | -20.71 | 5.07 |  |  |
| TG+WHR+FI | -18.89 | -15.27 |  |  |
| WHR | -25.22 | -1.00 |  |  |
| WHR+FI | -24.71 | -16.41 |  |  |

BMI, body mass index; TG, triglycerides; WHR, waist-to-hip ratio.

The univariate analyses showed that these signals would not have been detected in traditional EWAS for each trait separately, thus, indicating the improved power from the joint analysis of the correlated traits. We also characterized several established FG/FI or other relevant phenotype-associated EWAS signals through our MP-EWAS approach. A recent large-scale study in 4,808 (discovery) and 11,750 (replication) non-diabetic individuals by Liu *et al.* reported nine novel differentially methylated sites in whole blood with *P*<1.27×10^−7^: sites in *LETM1*, *RBM20*, *IRS2*, *MAN2A2* genes and 1q25.3 region were associated with FI; sites in *FCRL6*, *SLAMF1*, *APOBEC3H* genes and 15q26.1 region were associated with FG^22^. We show that of the sites associated with FI, 1q25.3 and *FAM92B* (not replicated in the original study) were also nominally (*P*<0.05) associated in our study, (**Table 4**). Similarly of the FG-associated sites, *BRE* (not replicated in the original study) was nominally associated in our study (**Table 4**).

**Table 4.** Sites reaching epigenome-wide significance in the study by Liu *et al.*^22^ and their associations in META-methylSCOPA meta-analysis within our study sample including 1,100 individuals from NFBC1966 and NFBC1986. Measured potential confounders body mass index (BMI), triglycerides (TG) and waist-to-hip ratio (WHR) were included in Model 1 whereas their effects were regressed out in Model 2.

| Locus | CpG | Chr | Position | Regulatory feature | Trait | Model 1 P-value | Model 2 P-value |
| --- | --- | --- | --- | --- | --- | --- | --- |
| *FCRL6*# | cg00936728 | 1 | 159772194 | NA | Glucose | 0.45 | 0.24 |
| *SLAMF1*# | cg18881723 | 1 | 160616870 | Promoter associated | Glucose | 0.74 | 0.68 |
| *1q25.3*# | cg13222915 | 1 | 184598594 | NA | Insulin | 0.014 | 0.24 |
| *BRE* | cg20657709 | 2 | 28509570 | NA | Glucose | 0.091 | 0.039 |
| *LRPPRC* | cg01913188 | 2 | 44223249 | Promoter associated | Glucose | 0.38 | 0.87 |
| *IRAK2* | cg14527942 | 3 | 10276383 | NA | Insulin | 0.31 | 0.69 |
| *LETM1*# | cg13729116 | 4 | 1859262 | Promoter associated | Insulin | 0.41 | 0.53 |
| *RBM20*# | cg15880704 | 10 | 112546110 | NA | Insulin | 0.52 | 0.61 |
| *IRS2*# | cg25924746 | 13 | 110432935 | Promoter associated | Insulin | 0.014 | 0.77 |
| *SPTB* | cg07119168 | 14 | 65225253 | NA | Glucose | 0.84 | 0.33 |
| *15q26.1*# | cg18247172 | 15 | 91370233 | NA | Glucose | 0.025 | 0.05 |
| *MAN2A2*# | cg20507228 | 15 | 91460071 | Promoter associated (Cell type specific) | Insulin | 0.13 | 0.10 |
| *FAM92B* | cg06709610 | 16 | 85143924 | NA | Insulin | 0.077 | 0.080 |
| *CD300A* | cg08087047 | 17 | 72461209 | NA | Glucose | 0.51 | 0.28 |
| *APOBEC3*# | cg06229674 | 22 | 39492189 | NA | Glucose | 0.63 | 0.77 |

#Replicated in Liu et al.

Another study by Zaghlool *et al*.^23^ aimed at elucidating the molecular pathways of 20 previously established CpG sites by using multi-omics data in 359 samples from the multi-ethnic Qatar Metabolomics Study on Diabetes. We observe associations at nominal significance at six of these 20 sites and demonstrate associations within Model 1 at *PHGDH, TXNIP, SLC7A11, CPT1A, MYO5C* and *ABCG1* through the dissection of multi-phenotype effects within our relatively small study sample (**Table 5**).

**Table 5.** Previously established sites for diabetes and its risk factors and their associations within our study sample including 1,100 individuals from NFBC1966 and NFBC1986. Measured potential confounders body mass index (BMI), triglycerides (TG) and waist-to-hip ratio (WHR) were included in Model 1 whereas their effects were regressed out in Model 2.

| Locus | Trait | CpG | Chr | Position | Model 1<br>*P*-value | Model 2<br>*P*-value |
| --- | --- | --- | --- | --- | --- | --- |
| *DHCR24* | Obesity | cg17901584 | 1 | 55353706 | 0.075 | 0.53 |
| *GFI1* | Smoking | cg09935388 | 1 | 92947588 |  |  |
| *PHGDH* | BMI, blood pressure, liver function | cg14476101 | 1 | 120255992 | 0.0056 | 0.031 |
| *TXNIP* | T2D | cg19693031 | 1 | 145441552 | 8.36E-04 | 0.051 |
|  | Metabolite, lipid | cg23079012 | 2 | 8343710 | 0.94 | 0.55 |
| *ALPPL2* | Smoking | cg21566642 | 2 | 233284661 | 0.25 | 0.18 |
| *UGT2B15* | Metabolite | cg09189601 | 4 | 69514031 | 0.56 | 0.19 |
| *SLC7A11* | Obesity | cg06690548 | 4 | 139162808 | 0.0037 | 0.44 |
| *AHRR* | Smoking | cg05575921 | 5 | 373378 | 0.40 | 0.12 |
|  | Metabolite, protein, smoking | cg06126421 | 6 | 30720080 | 0.26 | 0.45 |
| *LOC1001323546* | BMI, blood pressure | cg18120259 | 6 | 43894639 | 0.15 | 0.087 |
| *SLC25A22* | Metabolite, T2D, BMI | cg09441501 | 11 | 798350 | 0.56 | 0.67 |
| *CPT1A* | T2D and obesity | cg00574958 | 11 | 68607622 | 1.74E-07 | 0.91 |
|  |  | cg13526915 | 14 | 24164078 |  |  |
| *MYO5C* | Obesity | cg06192883 | 15 | 52554171 | 6.53E-04 | 0.48 |
| *TPM1* | Metabolite, smoking | cg10403394 | 15 | 63349192 | 0.32 | 0.85 |
| *RARA* | Smoking | cg19572487 | 17 | 38476024 | 0.72 | 0.93 |
| *F2RL3* | Smoking | cg03636183 | 19 | 17000585 | 0.42 | 0.56 |
| *SLC1A5* | BMI, blood pressure | cg22304262 | 19 | 47287778 | 0.33 | 0.87 |
| *ABCG1* | T2D and obesity | cg06500161 | 21 | 43656587 | 2.58E-06 | 0.077 |

## Conclusions

We have extended the multi-phenotype analyses to the EWAS framework and implemented this in the publicly available software tools methylSCOPA and METAmethylSCOPA. The application of the method to glycaemic traits demonstrated its enhanced power over single-trait EWAS for correlated phenotypes in large-scale data and the ability to characterize signals that are associated with correlated phenotypes.

## Availability and requirements

Project name: methylSCOPA.

Availability: the methylSCOPA and META-methylSCOPA tutorial can be found on “http://www.imperial.ac.uk/people/h.draisma/research.html”. The methylSCOPA and META-methylSCOPA software can be found on “http://doi.org/10.5281/zenodo.1137744” and on “https://doi.org/10.5281/zenodo.1286392”, respectively.

Operating system(s): Linux.

Programming language: C++ (including files from the ALGLIB project for statistical analysis and the TCLAP project for command line argument parsing).

Any restrictions on use by academics: none.

## Acknowledgments

Northern Finland Birth Cohort (NFBC1966) would like to thank the late professor Paula Rantakallio (launch of NFBC1966), the participants in the 31 year study and the NFBC project center.

## Availability of data and materials

The methylSCOPA and METAmethylSCOPA software tools are freely available at URL: xxx.

The Northern Finland Birth Cohort data which were used for the application of the developed tool are available upon collaboration and formal data request only, please see http://www.oulu.fi/nfbc/node/18136.

